# Host range expansion of *Shigella* phage Sf6 evolves through the dual roles of its tailspike

**DOI:** 10.1101/2022.04.06.487239

**Authors:** Sundharraman Subramanian, John A. Dover, Kristin N. Parent, Sarah M. Doore

**Affiliations:** Department of Biochemistry and Molecular Biology, Michigan State University, East Lansing, MI 48824, USA; Department of Microbiology and Cell Science, University of Florida, Gainesville, FL 36211, USA

## Abstract

The first critical step in a virus’s infection cycle is attachment to its host. This interaction is precise enough to ensure the virus will be able to productively infect the cell, but some flexibility can be beneficial to enable co-evolution and host range switching or expansion. Like many bacterial viruses, bacteriophage Sf6 utilizes a two-step process to recognize and attach to its host, *Shigella flexneri*. Sf6 first recognizes the lipopolysaccharide (LPS) structure of *S. flexneri*, then binds to either outer membrane protein (Omp) A or OmpC. This phage typically infects serotype Y strains but can also form small, turbid plaques on serotype 2a_2_ with greatly reduced plating efficiency, suggesting inefficient infection. To examine the interactions between Sf6 and this sub-optimal host, phage were experimentally evolved using mixed populations of *S. flexneri* serotypes Y and 2a_2_. The recovered mutants could infect serotype 2a_2_ with greater efficiency than the ancestral Sf6, forming clear plaques on both serotypes. All mutations mapped to two distinct regions of the tailspike protein: 1) adjacent to, but not part of, the LPS binding site near the N-terminus; and 2) at the distal, C-terminal tip of the protein. Rather than weak interactions between the Sf6 tailspike and 2a_2_ O-antigen, LPS of this serotype appears to inhibit infection by binding the wild-type particles more strongly, effectively removing them from the environment. These mutations reduce the inhibitory effect by either reducing electrostatic interactions with the O-antigen or increasing reliance on the Omp secondary receptors.

## Introduction

Bacteriophages, also known as phages, are viruses that infect bacteria. These viruses are diverse, ancient, and ubiquitous (Pope, et al. 2015). Phages persist in a variety of conditions, within fluctuating or stable environments, and likely encounter numerous different types of cells. Like all viruses, a fundamental step in the phage infection cycle is recognition of a host, including the ability to discriminate between susceptible and non-susceptible hosts. For many tailed phages infecting Gram-negative bacteria, this often involves a two-step process: first, reversible binding to a primary receptor; second, irreversible binding to a secondary receptor (Bertozzi Silva, et al. 2016; Subramanian, et al. 2020). The primary receptor is often the lipopolysaccharide (LPS) that extends from the cell surface and coats the membrane. The secondary receptor is more limited, such as one or two types of proteins embedded in the membrane. Using these mechanisms, bacteriophage host range can be broad, with a phage able to infect multiple species or genera; or narrow, with a phage being restricted to hosts with specific LPS chemistry.

Members of the genus *Shigella* are Gram-negative bacterial human pathogens causing bacillary dysentery, with infection usually linked to contaminated food or water sources (Zaidi and Estrada-Garcia 2014; Connor, et al. 2015). *Shigella* persists in environmental surfacewater, where it routinely emerges and infects over 250 million people annually, resulting in approximately 212 thousand deaths (Collaborators 2018). Once ingested, the bacteria proliferate intracellularly in epithelial cells of the large intestine, causing severe diarrhea and dehydration (Zaidi and Estrada-Garcia 2014; Anderson, et al. 2016; Killackey, et al. 2016). Of all *Shigella* species, *S. flexneri* is most frequently found in developing countries but has been responsible for outbreaks across the world, with different serotypes predominating different regions (Connor, et al. 2015; Anderson, et al. 2016). The serotype is determined by the repeating sugar units of the O-antigen and their subsequent glucosylation or acetylation, with Y being the simplest, least decorated type (Sun, et al. 2014; Muthuirulandi Sethuvel, et al. 2017). To date, over 50 O-antigen variants have been described for *S. flexneri*, with new serotypes and subtypes regularly emerging, often due to a combination of serotype-converting prophages and selection for virulence or immune evasion (Allison and Verma 2000; Knirel, et al. 2015; Muthuirulandi Sethuvel, et al. 2017). This property has made vaccine development particularly challenging (Levine, et al. 2007).

Since attachment and genome ejection are the first stages of interaction between virus and host, determining how phages recognize a given serotype is critical to our understanding of phage biology and ecology, as is how they may switch between serotypes or expand into new serotypes. *Shigella flexneri* phage Sf6 is a member of the *Podoviridae* family and is closely related to phage P22. Using cryo-electron tomography, the steps involved in the initial contact between P22 and its host *Salmonella enterica* sv. Typhimurium have been visualized (Wang, et al. 2019), and the structures of both P22 and Sf6 tail proteins have been resolved by various methods (Steinbacher, et al. 1996; Olia, et al. 2007; Muller, et al. 2008; Bhardwaj, et al. 2011; Parent, et al. 2012; Pintilie, et al. 2016; Wang, et al. 2019; Subramanian, et al. 2020). As part of the tail machinery, the tailspikes of P22 and Sf6 recognize and bind the O-antigen of LPS, then cleave the sugars using endorhamnosidase activity, which brings the phage closer to the host membrane (Figure 1) (Chua, et al. 1999; Muller, et al. 2008). In the case of Sf6, this lets the particle subsequently interact with outer membrane proteins (Omps) A or C as their secondary receptors (Parent, et al. 2014; Porcek and Parent 2015). In general, phage-mediated cleavage of LPS has been extensively studied, and the structures and catalytic mechanisms are known for a variety of tailspike homologs (Steinbacher, et al. 1994; Barbirz, et al. 2008; Muller, et al. 2008; Andres, et al. 2012; Lee, et al. 2017; Broeker, et al. 2019; Plattner, et al. 2019). It has also been determined that phage P22 interacts weakly with O-antigen derived from non-host *Salmonella enterica* sv. Paratyphi, with the oligosaccharide occupying an energetically unfavorable conformation (Steinbacher, et al. 1994; Barbirz, et al. 2008; Muller, et al. 2008; Andres, et al. 2012; Lee, et al. 2017; Broeker, et al. 2019; Plattner, et al. 2019). For P22, LPS alone is sufficient to induce genome ejection *in vitro*, suggesting tighter binding with this non-host LPS may confer broader host range (Andres, et al. 2010). However, Sf6 requires both LPS *and* a secondary receptor to eject its genome (Parent, et al. 2014; Porcek and Parent 2015). How the latter type of phage adapts to infect new hosts through interactions with both LPS and secondary receptors has not received much attention.

**Figure 1.**
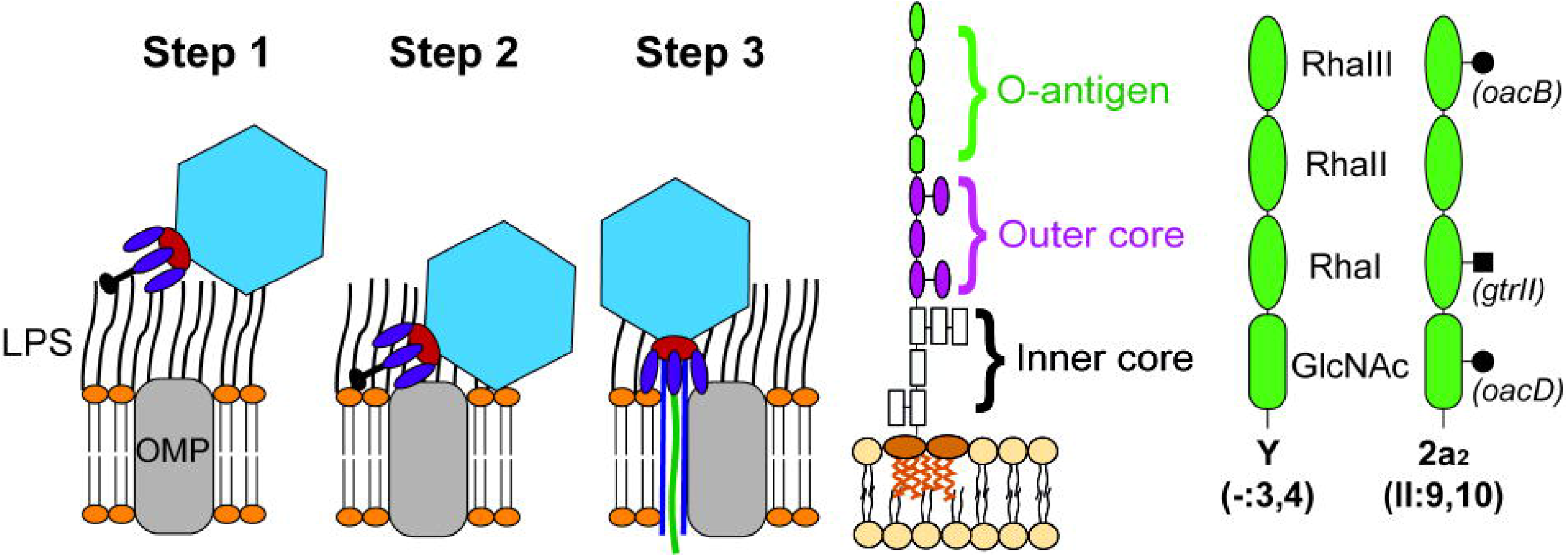
Overview of Sf6 virus attachment and infection (left), with the general structure of lipopolysaccharide (LPS) indicated in the center. Differences between the serotypes of host strains PE577 and CFS100, Y and 2a_2_ respectively, are shown (right). Modified from (Parent, et al. 2014; Porcek and Parent, 2015; Bohm, et al. 2018; Teh, et al. 2020).

With *S. flexneri* O-antigens constantly changing and new types frequently emerging, it seems crucial for phages that use O-antigen to either adapt to different serotypes or maintain a broader host range. Despite this, phage Sf6 infects a narrow range of *S. flexneri*, being primarily limited to serotype Y hosts (Gemski, et al. 1975). Previous experimental evolution results also suggest that another podovirus, T7, reproducibly contracts its host range when passaged in the presence of multiple non-susceptible hosts (Holtzman, et al. 2020), suggesting specialization may be more favorable in multi-species environments. In this study, we experimentally evolved phage Sf6 in the presence of both susceptible serotype Y and non-susceptible serotype 2a_2_ strains of *S. flexneri*, then isolated mutants with expanded host range. The O-antigen between serotypes Y and 2a_2_ differ by three modifications, as shown in Figure 1. To capture as many potential routes of evolution as possible, we employed a high-throughput method to screen approximately 300 individually passaged populations. Our results show that the tailspike protein is the critical structure for mediating not only LPS interactions, but also for identifying the secondary receptor, particularly OmpC in this scenario. In addition, phage-LPS interactions were previously thought of as reversible if the phage bound to a non-susceptible host. Here, we show that phage-LPS interactions can be detrimental to the phage. Rather than binding weakly or reversibly, bacterial LPS can bind in a way that inactivates the phage tail, rendering the particle no longer infectious. Similar to OMVs, LPS may contribute to “herd immunity” in a mixed community of bacteria and phage, but with reduced cost. A more thorough understanding of these interactions between *S. flexneri* and its phages will therefore lead to understanding the ecology and evolution of *S. flexneri* persistence in the environment and facilitate the development of tools for treating infections or detecting its presence.

## Results

### Mutations conferring broader host range map to the tailspike gene

Initially, we attempted to experimentally evolve Sf6 using single-step selections on bacterial lawns or serial passages in liquid culture with only the new serotype 2a_2_ host CFS100. These attempts with non-susceptible hosts alone were unsuccessful: no plaques were recovered, and all phage populations died out within the first few passages. To avoid this issue, our next trial in bulk liquid culture used a mixture of 90% non-susceptible host (serotype 2a_2_, CFS100) and 10% susceptible host (serotype Y, PE577). Using this method, we were able to isolate one mutant that could form plaques on CFS100. After whole genome sequencing, a single mutation was identified in the tailspike *gp14* gene, which substituted asparagine 455 to isoleucine (N455I). Afterwards, to increase our ability to recover and detect mutants with expanded host range, we used deep-well 96-well plates and 0.5 mL volumes to propagate phage in parallel (see Supplmentary Figure S1 for evolution scheme). At the end of each incubation period, the bacteria were lysed with chloroform so only the phage were passed to a fresh culture. Using this method, approximately 300 replicate populations were serially passaged, from which 34 isolates were found to infect the new host, CFS100. The recovered mutants were plaque purified to ensure each phage was isogenic, then amplified to high titer for subsequent analysis. Initial characterization included whole genome sequencing of purified genomic DNA, which revealed that each mutant had only 1 or 2 base pair (bp) changes in the entire ~40 kbp genome, which is consistent with previous Sf6 evolution schemes (Dover, et al. 2016). As before, all mutations were in the Sf6 *gp14* gene, which encodes the tailspike protein. This suggested these point mutations were necessary and sufficient to confer the phenotype, and that the tailspike is critical for determining host range. The tailspike protein is present as six trimers surrounding the portal and tail needle (represented by purple proteins in Figure 1). Each trimer exhibits endorhamnosidase activity, responsible for cleaving the O-antigen repeating unit of LPS, with key catalytic residues found on different subunits that line the groove between their adjacent β-helices (Figure 2, blue residues) (Muller, et al. 2008). Although 34 individual isolates were identified, these represent eight non-synonymous mutations and nine genotypes. On the tailspike protein, these changes cluster around two distinct regions: one below the LPS binding site and the other at the distal tip of the protein (Figure 2, pink and yellow residues). The changes highlighted in pink, A426G and N455I, arose the most frequently, representing 16 and 11 isolates respectively (see Supplementary Table S1 for a complete list of mutants isolated). These mutations were also found in tandem with other changes to produce four double mutants: A426G/N455I, A426G/N508T, A426G/T564R, and N455I/N556Y. The three remaining mutants were single non-synonymous mutations: T443P, G585D, or S589A.

**Figure 2.**
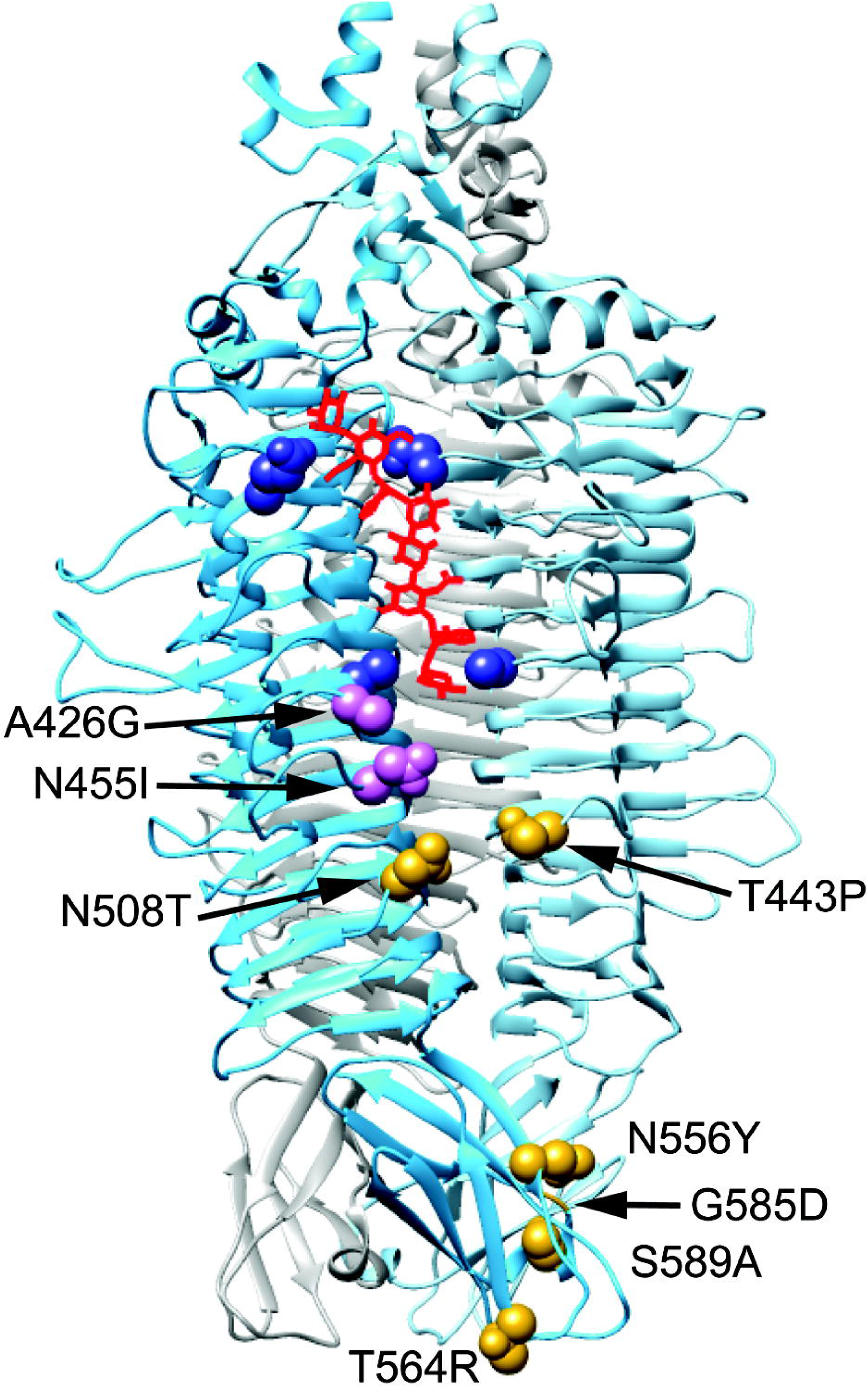
Crystal structure of the Sf6 tailspike trimer (PDB 2VBM, (Muller, et al. 2008)), with the location of recovered mutations mapping to the indicated sites. A model of LPS is shown in red, with catalytic residues highlighted in dark blue. Mutations recovered most frequently are in pink; all other mutations are in gold.

### Mutants are cold sensitive on the new host

To determine how efficiently these mutants could infect the 2a_2_ serotype host, and under which parameters, a series of quantitative plaque assays were conducted. First, mutants were examined at various temperatures on both the serotype Y and serotype 2a_2_ hosts. All except two mutants were completely unable to form plaques at or below 25 °C on CFS100 (Table 1). It was previously reported that *S. flexneri* produces greater levels of LPS and lower levels of OmpC at lower temperatures (Niu, et al. 2013). Since OmpC was previously identified as a secondary receptor for ancestral Sf6 (Parent, et al. 2014), we hypothesized that reduction of OmpC production at lower temperatures may have affected the plating efficiency for these tailspike mutants. The mechanism behind this cold sensitive (*cs*) phenotype was therefore investigated by making a series of genetic knockouts (see Supplementary Table S2 for a complete list of strains used). These knockouts targeted either the bacterial LPS structure, outer membrane proteins A or C, or a combination of both. For LPS, knockouts of *waaL* and *gtrII* affect the O-antigen by either removing it entirely, or by removing the glucosyl group of the 2a_2_ repeating unit, respectively.

**Table 1.** Efficiency of plating of ancestral Sf6 and mutants at different temperatures. Red shading indicates low efficiency of plating values (≤ 10^-2^).

| Phage Genotype | Serotype Y |  | Serotype 2a |  | Serotype 2a/*AgtrII* |  |
| --- | --- | --- | --- | --- | --- | --- |
|  | 25°C | 37°C | 25°C | 37°C | 25°C | 37°C |
| Ancestor | 0.3 ± 0.2 | 1.0 | < 10<sup>-6</sup> | < 10<sup>-6</sup> | 0.3 ± 0.0 | 0.9 ± 0.1 |
| A426G | 0.4 ± 0.2 | 1.0 | < 10<sup>-6</sup> | 0.6 ± 0.0 | 0.8 ± 0.2 | 0.7 ± 0.1 |
| A426G/N455I | 0.6 ± 0.3 | 1.0 | 0.1 ± 0.0 | 0.5 ± 0.1 | 0.5 ± 0.2 | 0.7 ± 0.2 |
| A426G/N508T | 0.5 ± 0.2 | 1.0 | < 10<sup>-6</sup> | 0.5 ± 0.1 | 0.9 ± 0.6 | 0.8 ± 0.1 |
| A426G/T564R | 0.5 ± 0.4 | 1.0 | < 10<sup>-6</sup> | 0.6 ± 0.1 | 0.9 ± 0.6 | 0.7 ± 0.1 |
| T443P | 0.6 ± 0.2 | 1.0 | < 10<sup>-6</sup> | 0.7 ± 0.1 | 0.6 ± 0.0 | 0.9 ± 0.1 |
| N455I | 0.2 ± 0.1 | 1.0 | < 10<sup>-6</sup> | 0.5 ± 0.1 | 0.8 ± 0.3 | 0.6 ± 0.1 |
| N455I/N556Y | 0.1 ± 0.0 | 1.0 | 0.1 ± 0.0 | 0.2 ± 0.1 | 1.0 ± 0.6 | 0.7 ± 0.2 |
| G585D | 0.1 ± 0.0 | 1.0 | < 10<sup>-6</sup> | 10<sup>-2</sup> | 1.1 ± 1.0 | 0.8 ± 0.2 |
| S589A | 0.5 ± 0.1 | 1.0 | < 10<sup>-6</sup> | 10<sup>-2</sup> | 0.9 ± 0.2 | 0.8 ± 0.2 |

As shown in Table 2, *ΔwaaL* prevents plaque formation of any phage isolate in either background, consistent with previous results showing that Sf6 requires O-antigen to attach to its host (Muller, et al. 2008). Removing *gtrII* in the CFS100 background restored the ability of all mutants to form plaques at all temperatures, including 25°C, suggesting this glucosyl group is necessary to confer resistance. Conversely, removing *ompC* reduced plaque formation on the new host for all mutants, though this phenotype was most severe for G585D and S589A. These two mutants showed reduced efficiency of plating in both CFS100/*ΔompA* and CFS100/*ΔompC* strains, suggesting the residues at the distal tip of the tailspike mediate interactions with these proteins. Even in the original serotype Y host, plaque formation of S589A is reduced 10-fold in the *ΔompC* strain, highlighting the importance of S589 in interactions with the secondary protein receptor. These electrostatic mutations are complementary and consistent with recent work identifying electrostatic residues in the surface loops of OmpC as being important for *Shigella* phage attachment (Tinney, et al. 2022). In addition, since the OmpC protein sequence is identical between these two strains, the difference in plating efficiency can be attributed to the differences in the LPS.

**Table 2.**
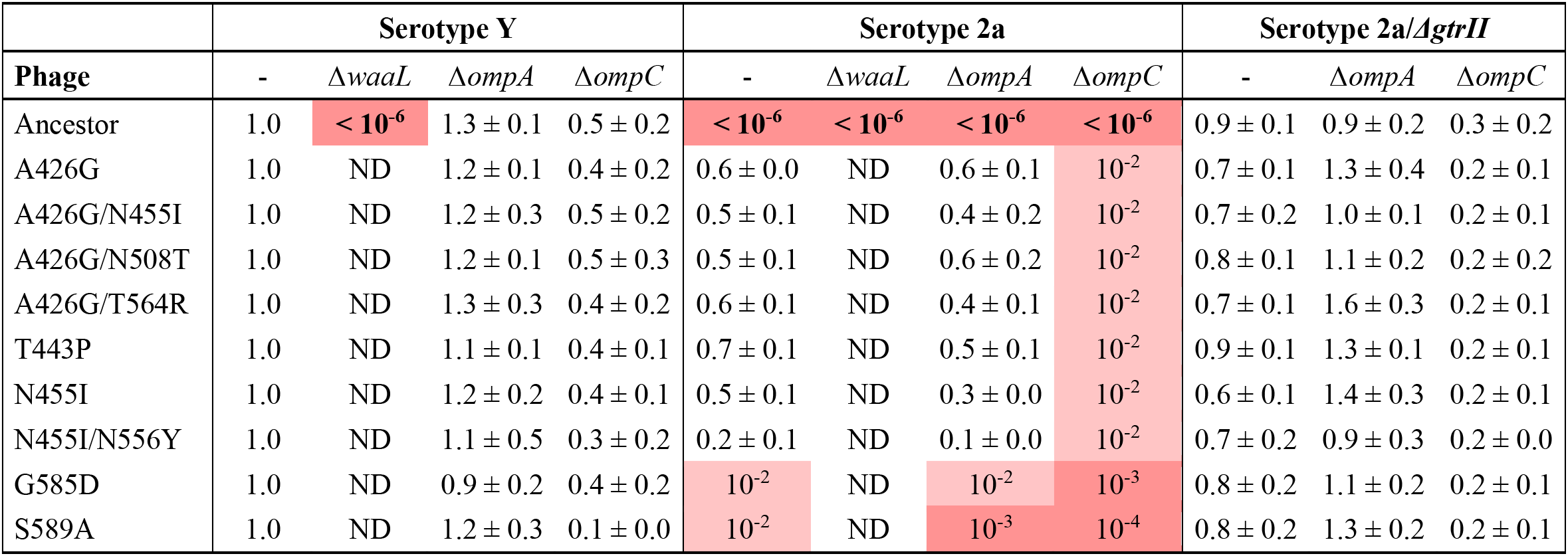
Efficiency of plating of ancestral Sf6 and mutants on strains with or without the indicated receptors at 37 °C. Red shading indicates low efficiency of plating values (≤ 10^-2^).

|  | Serotype Y |  |  |  | Serotype 2a |  |  |  | Serotype 2a/*AgtrII* |  |  |
| --- | --- | --- | --- | --- | --- | --- | --- | --- | --- | --- | --- |
| Phage | - | $\Delta waaL$ | $\Delta ompA$ | $\Delta ompC$ | - | $\Delta waaL$ | $\Delta ompA$ | $\Delta ompC$ | - | $\Delta ompA$ | $\Delta ompC$ |
| Ancestor | 1.0 | $< 10^{-6}$ | $1.3 \pm 0.1$ | $0.5 \pm 0.2$ | $< 10^{-6}$ | $< 10^{-6}$ | $< 10^{-6}$ | $< 10^{-6}$ | $0.9 \pm 0.1$ | $0.9 \pm 0.2$ | $0.3 \pm 0.2$ |
| A426G | 1.0 | ND | $1.2 \pm 0.1$ | $0.4 \pm 0.2$ | $0.6 \pm 0.0$ | ND | $0.6 \pm 0.1$ | $10^{-2}$ | $0.7 \pm 0.1$ | $1.3 \pm 0.4$ | $0.2 \pm 0.1$ |
| A426G/N455I | 1.0 | ND | $1.2 \pm 0.3$ | $0.5 \pm 0.2$ | $0.5 \pm 0.1$ | ND | $0.4 \pm 0.2$ | $10^{-2}$ | $0.7 \pm 0.2$ | $1.0 \pm 0.1$ | $0.2 \pm 0.1$ |
| A426G/N508T | 1.0 | ND | $1.2 \pm 0.1$ | $0.5 \pm 0.3$ | $0.5 \pm 0.1$ | ND | $0.6 \pm 0.2$ | $10^{-2}$ | $0.8 \pm 0.1$ | $1.1 \pm 0.2$ | $0.2 \pm 0.2$ |
| A426G/T564R | 1.0 | ND | $1.3 \pm 0.3$ | $0.4 \pm 0.2$ | $0.6 \pm 0.1$ | ND | $0.4 \pm 0.1$ | $10^{-2}$ | $0.7 \pm 0.1$ | $1.6 \pm 0.3$ | $0.2 \pm 0.1$ |
| T443P | 1.0 | ND | $1.1 \pm 0.1$ | $0.4 \pm 0.1$ | $0.7 \pm 0.1$ | ND | $0.5 \pm 0.1$ | $10^{-2}$ | $0.9 \pm 0.1$ | $1.3 \pm 0.1$ | $0.2 \pm 0.1$ |
| N455I | 1.0 | ND | $1.2 \pm 0.2$ | $0.4 \pm 0.1$ | $0.5 \pm 0.1$ | ND | $0.3 \pm 0.0$ | $10^{-2}$ | $0.6 \pm 0.1$ | $1.4 \pm 0.3$ | $0.2 \pm 0.1$ |
| N455I/N556Y | 1.0 | ND | $1.1 \pm 0.5$ | $0.3 \pm 0.2$ | $0.2 \pm 0.1$ | ND | $0.1 \pm 0.0$ | $10^{-2}$ | $0.7 \pm 0.2$ | $0.9 \pm 0.3$ | $0.2 \pm 0.0$ |
| G585D | 1.0 | ND | $0.9 \pm 0.2$ | $0.4 \pm 0.2$ | $10^{-2}$ | ND | $10^{-2}$ | $10^{-3}$ | $0.8 \pm 0.2$ | $1.1 \pm 0.2$ | $0.2 \pm 0.1$ |
| S589A | 1.0 | ND | $1.2 \pm 0.3$ | $0.1 \pm 0.0$ | $10^{-2}$ | ND | $10^{-3}$ | $10^{-4}$ | $0.8 \pm 0.2$ | $1.3 \pm 0.2$ | $0.2 \pm 0.1$ |

Many phages, including Sf6, identify and infect their host using a two-step process: a reversible attachment phase, followed by an irreversible phage during which the genome is ejected. To determine which of these two stages might be hindered in the wild-type phage in the presence of CFS100 and how the altered tailspikes could overcome this block to access a new host, both attachment and ejection were investigated. As shown in Figure 3A, nearly all wild-type phage particles attach to the serotype Y host by 5 minutes. The same result was observed for the serotype 2a_2_ host. Attachment kinetics for three mutants—A426G/N455I, N455I/N556Y, and G585D— were measured to determine whether attachment was affected during experimental evolution. None of the three mutants exhibited altered kinetics versus the ancestral isolate. Thus, the rate of attachment does not explain why Sf6 is unable to infect CFS100, nor does it appear to contribute to the expanded host range phenotype.

**Figure 3.**
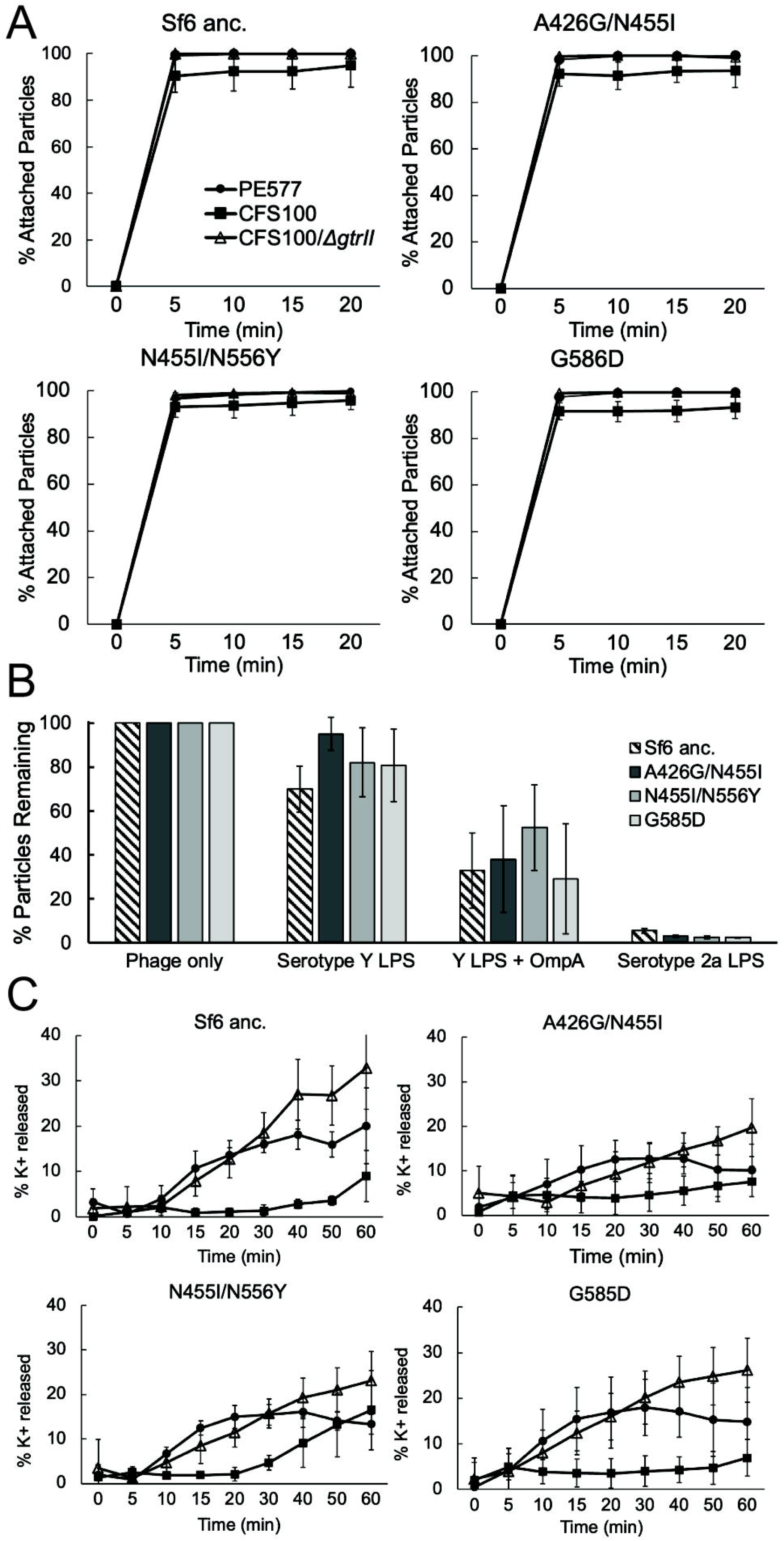
A) *In vivo* attachment kinetics, B) *in vitro* infectivity loss, and C) potassium efflux kinetics of ancestral Sf6 and three mutants.

### Incubation with serotype 2a_2_ LPS reduces Sf6 particle infectivity, but without genome ejection

Next, genome ejection was initially examined *in vitro* using purified components. Phage were incubated at 37°C alone, with LPS, the secondary receptor OmpA, or with both LPS and OmpA. Plaque assays were then used to measure remaining plaque forming units after the incubation period. Using this method, previous studies demonstrated that wild-type Sf6 requires both LPS and OmpA to ejects its genome *in vitro*, with LPS or OmpA alone being insufficient (Parent, et al. 2014; Porcek and Parent 2015). As expected, when incubated with serotype Y LPS alone, neither wild-type Sf6 nor the mutants showed significant loss of infectivity (Figure 3B). Conversely, adding both serotype Y LPS and OmpA resulted in most particles losing infectivity, consistent with genome ejection observed in (Porcek and Parent 2015). Surprisingly, when particles were incubated with serotype 2a_2_ LPS *alone*, this component was sufficient to significantly reduce particle infectivity. As with our attachment assay, this result was observed for both the ancestral and experimentally evolved Sf6 isolates.

Infectivity loss could be explained by one of two phenomena: 1) *in vitro* genome ejection occurred, meaning the now empty phage particles are unable to infect subsequent host cells due to genome loss; or 2) inhibition occurred, so the phage tail is blocked from binding to subsequent host cells, with no genome loss. To differentiate between these two possibilities, genome ejection was measured by potassium efflux. Upon penetration of the membrane and movement of the phage genome into the cell, potassium ions are released from the host and flow into the surrounding media: thus, measuring the change in potassium concentration serves as a more direct measure of genome ejection.

Ancestral Sf6 was incubated with serotype Y or serotype 2a_2_ host cells, and the change in potassium concentration was measured. As shown in Figure 3C, incubation with serotype Y host cells induced greater potassium release than serotype 2a_2_ host cells. This suggests that even by 60 minutes, at which point most particles lose their infectivity when incubated with serotype 2a LPS, very few particles are ejecting their genomes in the presence of this host.

### Serotype 2a_2_ lipopolysaccharide appears to render Sf6 non-infectious by sticking to tailspikes

Since both ancestral and evolved Sf6 isolates showed reduced infectivity without genome ejection in the presence of LPS derived from the serotype 2a_2_ host, we hypothesized that the non-host LPS was binding to phage particles more frequently or more strongly, resulting in tail protein blockage. To evaluate this possibility, the interactions between phage particles incubated with or without LPS were visualized directly. Aliquots from the *in vitro* infectivity loss assays were vitrified and examined by cryo-electron microscopy (cryoEM). Ancestral Sf6 and three mutants A426G/N455I, N455I/N556Y, and G585D were characterized under four conditions: phage alone, phage incubated at with serotype Y LPS derived from PE577, phage incubated with serotype 2a_2_ LPS derived from CFS100, or phage incubated with serotype Y_2_ LPS derived from CFS100/*ΔgtrII*. For each condition, approximately 500 particles were counted and scored as having: 1) an empty capsid, suggesting the genome had been ejected; 2) a full capsid, unbound to LPS; or 3) a full capsid, bound to LPS. Representative images of each condition are shown in Figure 4A. For ancestral Sf6, particles incubated with serotype Y LPS were primarily in the “free and full” category, regardless of whether the LPS was derived from PE577 or CFS100/*ΔgtrII*. Conversely, when phage were incubated with serotype 2a_2_ LPS, approximately 75% of the population were in the “bound and full” category. Combined with the potassium efflux data, this suggests the loss of infectivity is due to the phage tail proteins being stuck to LPS, rather than the LPS inducing genome ejection.

**Figure 4.**
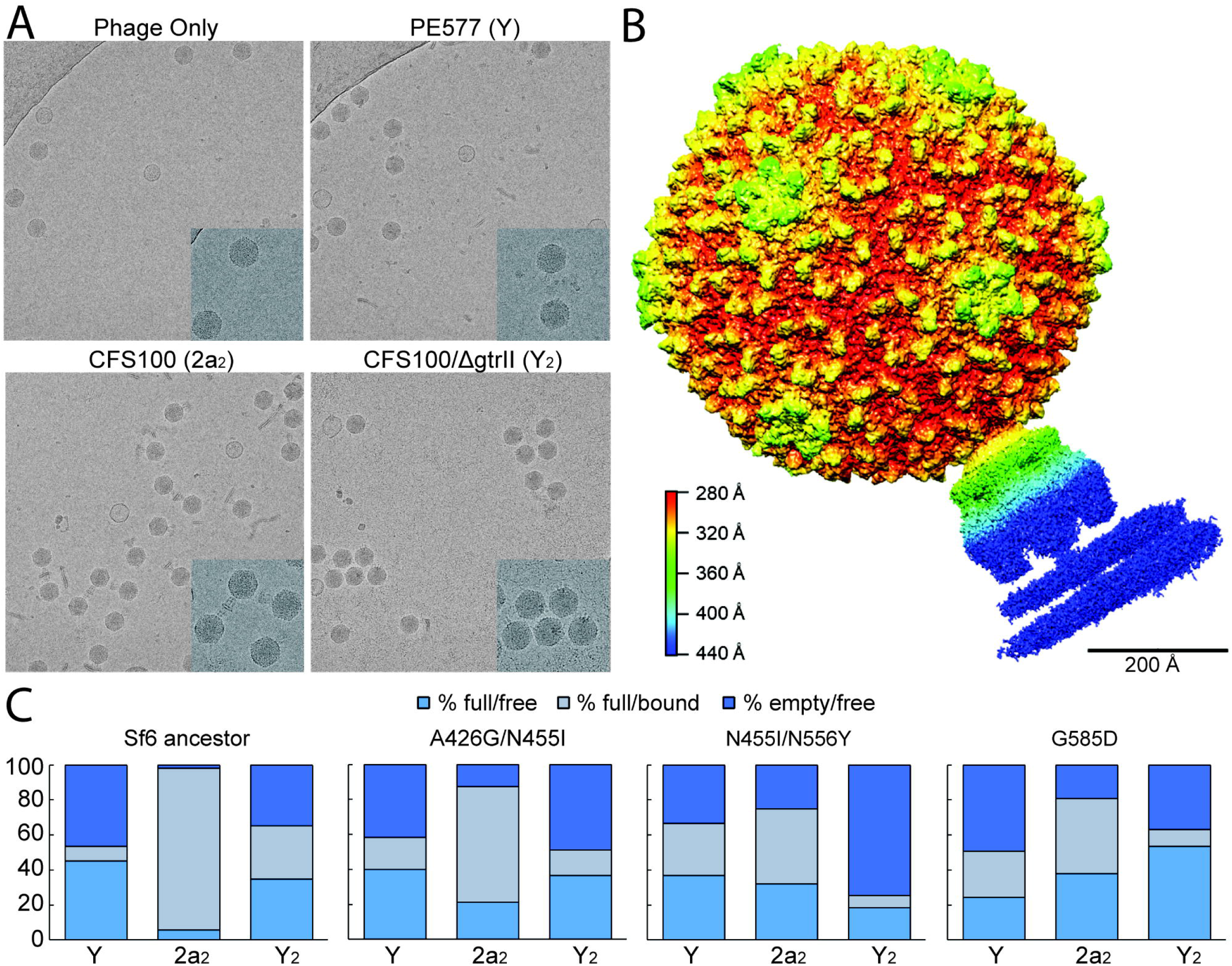
A) Representative images of the Sf6 ancestral strain imaged by cryoEM, either without LPS, or with Y, 2a_2_, or Y_2_ types of LPS; insets show greater detail of particles; B) Asymmetric reconstruction of Sf6 particles bound to serotype 2a LPS at 5.1 Å colored by radial distance; C) Quantitative results of particle status for Sf6 ancestral strain and three mutants based on images similar to those shown in A.

Although our results indicated a block early in the infection process, the specific stage in the sequence of attachment and infection kinetics was unclear. Previous studies of the closely-related phage P22 determined the series of events involved in phage attachment and genome ejection (Wang, et al. 2019). Using cryo-electron tomography (cryoET), the authors showed that the phage particles initially bind the host at an angle, with two adjacent tailspikes bound to LPS and the tail needle in contact with the cell surface. This observed orientation of phage P22 matches our Sf6 mutant data which shows that the OmpC binding region is in the distal tail tip, where Sf6 is likely interacting with the cell surface in a similar fashion (see Figure 1). The phage tailspikes then hydrolyze the O-antigen through a series of rapid release and rebinding events, bringing the rest of the tail closer to the cell surface. This is then followed by a reorientation of the particle, which positions the tail needle perpendicular to the membrane. The tail needle is then lost and the particle undergoes a dramatic rearrangement, which involves ejection proteins being released to form a tube. The genomic dsDNA is then translocated through this tube and into the host cell. The transition from oblique to perpendicular is inherently difficult to visualize but may involve additional tailspikes binding to and hydrolyzing nearby O-antigen molecules.

To determine how serotype 2a_2_ LPS may be inhibiting Sf6 particle infectivity, we performed an asymmetric reconstruction of the ancestral Sf6 strain bound to 2a_2_ LPS. If particles were stuck in the oblique orientation, we would expect to obtain a structure with the Sf6 tailspike bound at an angle, with contact between two tailspike trimers and the O-antigen of LPS. Conversely, if particles became stuck later, after reorientation, we would expect to see a perpendicular orientation, with the tail needle either present or absent (Wang, et al. 2019). This orientation was observed in a study by McNulty et al., using a mutant of phage P22 that lacked the tail needle protein (McNulty, et al. 2018). As shown in Figure 4B, the particle appears to be oriented perpendicular to the LPS, with the tail needle intact and the particles retaining their genomes. Although this experiment was performed using purified components rather than whole cells, this may suggest LPS can trap the particle just before the tail needle is lost, possibly with all six tailspike trimers bound to the O-antigen of LPS.

For the mutants, although most particles were still in the “full and bound” category when incubated with the serotype 2a LPS, a significantly greater proportion were free from LPS (Figure 4C). The three mutants analyzed also appeared to bind more frequently with serotype Y LPS when compared to ancestral Sf6, but this phenomenon did not transfer to the CFS100/Δ*gtrII* serotype Y_2_ LPS. Instead, in this last condition, the two double mutants A426G/N455I and N455I/N556Y were primarily empty particles, which may suggest altered genome ejection dynamics or stability in the presence of this type of Y_2_ LPS.

## Discussion

Bacteriophage Sf6 undergoes a two-step attachment process to identify its host: an initial reversible attachment to its primary receptor, LPS; then a second, irreversible attachment to its secondary receptor, OmpA or OmpC. Once irreversibly attached, the particle undergoes a rapid and dtrastic conformational change to translocate its genome across the membrane and into the host cell. The Sf6 tail complex is involved in each of these stages, and the tailspike is known to be critical for LPS binding and hydrolysis. The protein involved in irreversible attachment, however, remained unclear.

Here, we determined that the Sf6 gp14 tailspike is responsible for mediating multiple interactions between both LPS and the secondary receptors OmpA and OmpC. With one protein binding to both receptors, single point mutations in the *gp14* gene are sufficient to expand the Sf6 host range via altered interactions with the primary receptor, the secondary receptor, or both. Mutations near the LPS binding pocket may affect the strength of binding without affecting enzymatic activity of the protein. During preparation of this manuscript, Teh et al. also isolated an Sf6 mutant that could infect the serotype 2a_2_ host (Teh, et al. 2022). This mutant had three mutations in the tailspike: Q325L, A426G, and N508T. Two of these mutations, A426G and N508T, were also isolated here. The authors had previously shown that even the ancestral Sf6 could hydrolyze a minimal amount of serotype 2a_2_ LPS (~20-30% (Teh, et al. 2020; Teh, et al. 2022)), while the triple mutant could hydrolyze 68%. Rather than increasing binding affinity for 2a_2_ O-antigen, we hypothesize the mutations near the LPS binding site are either decreasing constraint of or altering electrostatic interactions with the O-antigen. These changes could allow for a better fit of the glucosylated 2a_2_ O-antigen structure, which likely assumes a helical rather than a linear shape (West, et al. 2005). Previous work using molecular dynamics demonstrated that mutations near the O-antigen binding site of the Sf6 tailspike can dramatically affect the flexibility of the site (Kunstmann, et al. 2020). As part of their study, Kunstmann et al. modeled a T443C substitution. While we isolated T443P here, the effect of increasing flexibility may be similar. Alternatively, by altering electrostatic interactions, the O-antigen may be less likely to become trapped in the binding pocket after hydrolysis, effectively reducing the “stickiness” of this type of LPS.

While many mutants appeared to overcome the host barrier via altered LPS interactions, others showed a much greater reliance on the secondary receptors OmpA and/or OmpC. These mutations, which all mapped to the distal end of the tailspike, may increase the binding affinity for these secondary receptors as a way to compensate for slow or stalled LPS hydrolysis. The G585D and S589A mutants had reduced plating efficiency on the CFS100 host compared to all other single or double mutants, suggesting this mechanism—while effective—may not be the most efficient.

Previous studies with closely-related phage P22 demonstrate that the phage binds certain non-host LPS weakly, allowing virions to detach from non-host cells (Andres, et al. 2013). By contrast, Sf6 appears to bind the serotype 2a_2_ non-host LPS strongly, with virions less able to release from nonhost cells. It has also been shown that *S. flexneri*, like other enteric bacteria, alters LPS and/or Omp production depending on environmental conditions, such as temperature (Maurelli, et al. 1984; Niu, et al. 2013). In freshwater reservoirs where *S. flexneri* persists, lower temperature would contribute to high levels of LPS production with low OmpC expression. In this case, phage may be more easily sequestered by LPS shed from the outer membrane of non-host cells, with a lower chance of overcoming the LPS barrier through secondary receptor binding. This mechanism of phage sequestration may provide a community benefit for bacteria, where susceptible and resistant strains may co-exist. Conversely, in the warmer environment of the human intestine where *S. flexneri* causes disease, phage may be able to overcome these barriers due to lower LPS production and/or greater OmpC expression.

In the arms race between bacteria and phages, many well-studied mechanisms occur at the cell surface. These include modifications to the cell receptor or phage receptor-binding proteins (Porcek and Parent 2015; Dunne, et al. 2019), reduced expression or shielding of receptors (van der Ley, et al. 1986; Meyer, et al. 2012; Tan, et al. 2015), and highly variable or horizontally transferred tail genes (Trojet, et al. 2011; Tzipilevich, et al. 2017). Outer membrane vesicles can also serve as decoys for bacteriophage (Manning and Kuehn 2011; Reyes-Robles, et al. 2018). Our results suggest that shedding LPS alone may also be sufficient to inactive bacteriophage in the environment, which would come at a greatly reduced resource cost. Using decoys rather than modifying receptor protein expression or structure is a method of herd immunity for bacteria. This mechanism can also reduce the size of the bacteriophage adaptive landscape by removing phage from the population. Since these decoys can sequester phage, it is important to determine how and when these decoys are most effective, and how phage can overcome this type of “passive” defense. In addition to understanding bacteria and bacteriophage ecology and evolution, it may be used to predict or inform the effectiveness of phage application for therapeutic use.

## Materials & Methods

### Bacterial strains and plasmids

The strains of bacteria used as hosts for Sf6, both PE577 and CFS100, are avirulent derivatives of *S. flexneri* and have been described (Morona, et al. 1994; Marman, et al. 2014; Doore, et al. 2021). Mutant strains including *ompA::kan, ompC::kan, gtrII::kan*, or a combination thereof were generated using previously-described methods (Doore, et al. 2021) based on lambda red recombineering (Datsenko and Wanner 2000). For double mutants, the first kanamycin cassette was removed by transforming the strain with pCP20, which harbors Flp recombinase. After recovery, bacteria were incubated overnight with ampicillin at 30 °C, then single colonies were streaked onto a new plate. The next day, single colonies were chosen and incubated at 42 °C to remove pCP20, then screened for loss of the kanamycin cassette by PCR.

### Phage methods and experimental evolution scheme

The ancestral isolate of Sf6 for this study is an obligately lytic mutant described in (Casjens, et al. 2004; Datsenko and Wanner 2000). Since the ability of Sf6 to infect CFS100 was very low (<10^-6^), we used a mixture of PE577 and CFS100 host strains during experimental evolution. Overnight cultures were therefore mixed in a ratio of 1:9 susceptible:non-susceptible strains to maintain selective pressure while allowing the phage to replicate in a restricted number of host cells. To maximize our ability to screen replicate populations, a deep-well 96-well plate was used with 0.5 mL total culture volume. At the beginning of each experiment, wells contained 450 μL Luria broth (LB), 30 μL of the 1:9 cell mixture, and 20 μL of a 10^6^ dilution of purified, isogenic Sf6, with a starting multiplicity of infection of ~0.1 phage per cell. This was then incubated at 37 °C while shaking for 4 hours for passage one. At the end of the incubation period, bacteria were lysed with chloroform and a replica pin tool was used to sequentially stamp 150×15 mm Petri plates seeded with non-susceptible CFS100 and then the susceptible PE577 host cells. Debris were then allowed to settle in the 96-well plates and the lysates were stored at 4°C overnight. The following morning, this process was repeated using 20 μL of lysate from the previous day rather than the isogenic starting phage stock and fresh cultures of the host cells retaining the 1:9 ratio. If a clearing was observed around a pin stamp on the CFS100 screening plate, the experiment was stopped and the phage from the corresponding well was plaque purified.

Once a clearing was observed on CFS100, the phenotype was confirmed by plaque purifying on CFS100 at least three times. A single plaque was then picked and amplified in CFS100 to create an isogenic stock. Once cleared, the culture was centrifuged for 20 min at 6 000 x g to remove cellular debris, then the supernatant was centrifuged for 90 min at 26 000 x g to pellet the phage. This pellet was then resuspended in phage dilution buffer (10mM Tris, 10mM MgCl2) by nutating at 4°C overnight. A final spin of 10 min at 6 000 x g removed any remaining cell debris, and the supernatant was retained as a high-titer phage stock.

To determine the genotype of mutant phages that could infect CFS100, genomic DNA was extracted from 1×10^10^-1×10^11^ phage particles using phenol-chloroform and sequenced at the Michigan State University Research Technology Support Facility (RTSF). Sequences were aligned using Breseq as previously described (Dover, et al. 2016).

### Attachment kinetics and in vitro infectivity loss assays

Attachment assays were preformed as has been described (Parent, et al. 2014). Briefly, phage were added to bacteria at an MOI of 10, then incubated at 37°C. Aliquots of the mixture were taken at the timepoints indicated and centrifuged for 30 sec to remove bacteria and any attached bacteriophages. The phages remaining in the supernatant were then quantified by plaque assay.

Infectivity loss assays were performed similar to *in vitro* genome ejection assays as previously described (Porcek and Parent 2015). Briefly, phage were incubated for 60 min in phage dilution buffer at 37 °C either alone, with purified LPS, or with purified LPS and OmpA. At the end of the incubation time, the mixture was serially diluted and infectious phage particles were quantified by plaque assay. After this additional incubation period, the mixture was again serially diluted, with infectious phage particles quantified by plaque assay.

### Cryo-electron microscopy and data processing

Small (3-5 μL) aliquots of virus/LPS mixtures were applied to R2/2 Quantifoil grids (Electron Microscopy Solutions) that had been glow discharged for 45 seconds in a Pelco Easiglow glow discharging unit. The samples were plunge frozen in liquid ethane using a Vitrobot Mark IV operated at 4°C and 100% humidity, with a blot force of 1 and 5 seconds of blotting time per grid.

For all virus/LPS mixtures, cryo-EM data were collected at the RTSF Cryo-EM facility using a Talos Arctica equipped with a Falcon 3 direct electron detector, operating at 200 keV under low dose conditions. Micrographs were collected at 45,000X nominal magnification (2.24 Å/pixel) by recording 11 frames over 3 sec for a total dose of 25 e^-^/Å^2^. For the Sf6-CFS100 LPS structure, cryo-EM data Sf6-CFS100 LPS were collected at Purdue Cryo-EM facility using a Titan Krios equipped with a K3 direct electron detector, and operating at 300 keV with a post-column GIF (20 eV slit width) under low dose conditions. Micrographs were collected at 53,000X nominal magnification (0.816 Å/pixel) by recording 40 frames over 4.4 sec for a total dose of 33 e^-^/Å^2^.

Dose-fractionated movies collected using the Falcon 3 direct electron detector were subjected to motion correction using the program MotionCor2. The resulting images were used for quantitative analysis of phage particles. Asymmetric reconstruction of Sf6 bound to CFS100 LPS was carried out using Relion 3.1.1. Briefly, the dose-fractionated movies were subjected to motion correction using Relion’s own implementation of MotionCor2. CTF estimation of the resulting images were estimated using CTFFIND-4.1 and particles were picked using the Autopick option. 4X binned particles were then extracted and subjected to 2D classification. Full cryoEM image processing statistics are listed in Supplementary Table S3. A total of 18,964 particles were used for 3D refinement, with the Sf6 virion map (EMD:5730, scaled by a factor of 0.65 in EMAN2, Symmetrized using Relion 3.0.8) serving as an initial model. Refined particles were extracted again with 2X binning and subjected to 3D refinement. The overall resolution was estimated based on the gold-standard Fourier shell correlation (FSC) = 0.143 criterion in the Post-process job. The final maps were deposited into EMDB (accession number EMD-26561).

## Acknowledgements

We would like to thank Jacob Taylor for preliminary work that helped develop our final evolutionary scheme. We would also like to thank Dr. Ian Molineux at the University of Texas – Austin for numerous helpful discussions. This work was supported by the National Institutes of Health GM110185, GM140803 and National Science Foundation CAREER Award 1750125 to K.N.P. and by startup funds from the University of Florida to S.M.D. We would like to thank the MSU RTSF Cryo-EM Core Facility for use of the Talos Arctica. Additionally, we thank Dr. T. Klose at Purdue University’s Midwest Cryo-EM Consortium (NIH Consortium #U24GM116789-03).

## References

Allison GE, Verma NK. 2000. Serotype-converting bacteriophages and O-antigen modification in Shigella flexneri. Trends Microbiol 8:17–23.

Anderson M, Sansonetti PJ, Marteyn BS. 2016. Shigella Diversity and Changing Landscape: Insights for the Twenty-First Century. Front Cell Infect Microbiol 6:45.

Andres D, Hanke C, Baxa U, Seul A, Barbirz S, Seckler R. 2010. Tailspike interactions with lipopolysaccharide effect DNA ejection from phage P22 particles in vitro. J Biol Chem 285:36768–36775.

Andres D, Roske Y, Doering C, Heinemann U, Seckler R, Barbirz S. 2012. Tail morphology controls DNA release in two Salmonella phages with one lipopolysaccharide receptor recognition system. Mol Microbiol 83:1244–1253.

Andres D, Gohlke U, Broeker NK, Schulze S, Rabsch W, Heinemann U, Barbirz S, Seckler R. 2013. An essential serotype recognition pocket on phage P22 tailspike protein forces Salmonella enterica serovar Paratyphi A O-antigen fragments to bind as nonsolution conformers. Glycobiology 23:486–494.

Barbirz S, Muller JJ, Uetrecht C, Clark AJ, Heinemann U, Seckler R. 2008. Crystal structure of Escherichia coli phage HK620 tailspike: podoviral tailspike endoglycosidase modules are evolutionarily related. Mol Microbiol 69:303–316.

Bertozzi Silva J, Storms Z, Sauvageau D. 2016. Host receptors for bacteriophage adsorption. FEMS Microbiol Lett 363.

Bhardwaj A, Molineux IJ, Casjens SR, Cingolani G. 2011. Atomic structure of bacteriophage Sf6 tail needle knob. J Biol Chem 286:30867–30877.

Bohm K, Porwollik S, Chu W, Dover JA, Gilcrease EB, Casjens SR, McClelland M, Parent KN. 2018. Genes affecting progression of bacteriophage P22 infection in Salmonella identified by transposon and single gene deletion screens. Mol Microbiol 108:288–305.

Broeker NK, Roske Y, Valleriani A, Stephan MS, Andres D, Koetz J, Heinemann U, Barbirz S. 2019. Time-resolved DNA release from an O-antigen-specific Salmonella bacteriophage with a contractile tail. J Biol Chem 294:11751–11761.

Casjens S, Winn-Stapley DA, Gilcrease EB, Morona R, Kuhlewein C, Chua JE, Manning PA, Inwood W, Clark AJ. 2004. The chromosome of Shigella flexneri bacteriophage Sf6: complete nucleotide sequence, genetic mosaicism, and DNA packaging. J Mol Biol 339:379–394.

Chua JEH, Manning PA, Morona R. 1999. The Shigella flexneri bacteriophage Sf6 tailspike protein (TSP)/endorhamnosidase is related to the bacteriophage P22 TSP and has a motif common to exo- and endoglycanases, and C-5 epimerases. Microbiology (Reading) 145 (Pt 7):1649–1659.

Collaborators GBDDD. 2018. Estimates of the global, regional, and national morbidity, mortality, and aetiologies of diarrhoea in 195 countries: a systematic analysis for the Global Burden of Disease Study 2016. Lancet Infect Dis 18:1211–1228.

Connor TR, Barker CR, Baker KS, Weill FX, Talukder KA, Smith AM, Baker S, Gouali M, Pham Thanh D, Jahan Azmi I, et al. 2015. Species-wide whole genome sequencing reveals historical global spread and recent local persistence in Shigella flexneri. Elife 4:e07335.

Datsenko KA, Wanner BL. 2000. One-step inactivation of chromosomal genes in Escherichia coli K-12 using PCR products. Proc Natl Acad Sci U S A 97:6640–6645.

Doore SM, Subramanian S, Tefft NM, Morona R, TerAvest MA, Parent KN. 2021. Large metabolic rewiring from small genomic changes between strains of Shigella flexneri. J Bacteriol.

Dover JA, Burmeister AR, Molineux IJ, Parent KN. 2016. Evolved Populations of Shigella flexneri Phage Sf6 Acquire Large Deletions, Altered Genomic Architecture, and Faster Life Cycles. Genome Biol Evol 8:2827–2840.

Dunne M, Rupf B, Tala M, Qabrati X, Ernst P, Shen Y, Sumrall E, Heeb L, Pluckthun A, Loessner MJ, et al. 2019. Reprogramming Bacteriophage Host Range through Structure-Guided Design of Chimeric Receptor Binding Proteins. Cell Rep 29:1336–1350 e1334.

Gemski P, Jr., Koeltzow DE, Formal SB. 1975. Phage conversion of Shigella flexneri group antigens. Infect Immun 11:685–691.

Holtzman T, Globus R, Molshanski-Mor S, Ben-Shem A, Yosef I, Qimron U. 2020. A continuous evolution system for contracting the host range of bacteriophage T7. Sci Rep 10:307.

Killackey SA, Sorbara MT, Girardin SE. 2016. Cellular Aspects of Shigella Pathogenesis: Focus on the Manipulation of Host Cell Processes. Front Cell Infect Microbiol 6:38.

Knirel YA, Sun Q, Senchenkova SN, Perepelov AV, Shashkov AS, Xu J. 2015. O-antigen modifications providing antigenic diversity of Shigella flexneri and underlying genetic mechanisms. Biochemistry (Mosc) 80:901–914.

Kunstmann S, Engstrom O, Wehle M, Widmalm G, Santer M, Barbirz S. 2020. Increasing the Affinity of an O-Antigen Polysaccharide Binding Site in Shigella flexneri Bacteriophage Sf6 Tailspike Protein. Chemistry 26:7263–7273.

Lee IM, Tu IF, Yang FL, Ko TP, Liao JH, Lin NT, Wu CY, Ren CT, Wang AH, Chang CM, et al. 2017. Structural basis for fragmenting the exopolysaccharide of Acinetobacter baumannii by bacteriophage PhiAB6 tailspike protein. Sci Rep 7:42711.

Levine MM, Kotloff KL, Barry EM, Pasetti MF, Sztein MB. 2007. Clinical trials of Shigella vaccines: two steps forward and one step back on a long, hard road. Nat Rev Microbiol 5:540–553.

Manning AJ, Kuehn MJ. 2011. Contribution of bacterial outer membrane vesicles to innate bacterial defense. BMC Microbiol 11:258.

Marman HE, Mey AR, Payne SM. 2014. Elongation factor P and modifying enzyme PoxA are necessary for virulence of Shigella flexneri. Infect Immun 82:3612–3621.

Maurelli AT, Blackmon B, Curtiss R, 3rd. 1984. Temperature-dependent expression of virulence genes in Shigella species. Infect Immun 43:195–201.

McNulty R, Cardone G, Gilcrease EB, Baker TS, Casjens SR, Johnson JE. 2018. Cryo-EM Elucidation of the Structure of Bacteriophage P22 Virions after Genome Release. Biophys J 114:1295–1301.

Meyer JR, Dobias DT, Weitz JS, Barrick JE, Quick RT, Lenski RE. 2012. Repeatability and contingency in the evolution of a key innovation in phage lambda. Science 335:428–432.

Morona R, Mavris M, Fallarino A, Manning PA. 1994. Characterization of the rfc region of Shigella flexneri. J Bacteriol 176:733–747.

Muller JJ, Barbirz S, Heinle K, Freiberg A, Seckler R, Heinemann U. 2008. An intersubunit active site between supercoiled parallel beta helices in the trimeric tailspike endorhamnosidase of Shigella flexneri Phage Sf6. Structure 16:766–775.

Muthuirulandi Sethuvel DP, Devanga Ragupathi NK, Anandan S, Veeraraghavan B. 2017. Update on: Shigella new serogroups/serotypes and their antimicrobial resistance. Lett Appl Microbiol 64:8–18.

Niu C, Shang N, Liao X, Feng E, Liu X, Wang D, Wang J, Huang P, Hua Y, Zhu L, et al. 2013. Analysis of Soluble protein complexes in Shigella flexneri reveals the influence of temperature on the amount of lipopolysaccharide. Mol Cell Proteomics 12:1250–1258.

Olia AS, Casjens S, Cingolani G. 2007. Structure of phage P22 cell envelope-penetrating needle. Nat Struct Mol Biol 14:1221–1226.

Parent KN, Erb ML, Cardone G, Nguyen K, Gilcrease EB, Porcek NB, Pogliano J, Baker TS, Casjens SR. 2014. OmpA and OmpC are critical host factors for bacteriophage Sf6 entry in Shigella. Mol Microbiol 92:47–60.

Parent KN, Gilcrease EB, Casjens SR, Baker TS. 2012. Structural evolution of the P22-like phages: comparison of Sf6 and P22 procapsid and virion architectures. Virology 427:177–188.

Pintilie G, Chen DH, Haase-Pettingell CA, King JA, Chiu W. 2016. Resolution and Probabilistic Models of Components in CryoEM Maps of Mature P22 Bacteriophage. Biophys J 110:827–839.

Plattner M, Shneider MM, Arbatsky NP, Shashkov AS, Chizhov AO, Nazarov S, Prokhorov NS, Taylor NMI, Buth SA, Gambino M, et al. 2019. Structure and Function of the Branched Receptor-Binding Complex of Bacteriophage CBA120. J Mol Biol 431:3718–3739.

Pope WH, Bowman CA, Russell DA, Jacobs-Sera D, Asai DJ, Cresawn SG, Jacobs WR, Hendrix RW, Lawrence JG, Hatfull GF, et al. 2015. Whole genome comparison of a large collection of mycobacteriophages reveals a continuum of phage genetic diversity. Elife 4:e06416.

Porcek NB, Parent KN. 2015. Key residues of S. flexneri OmpA mediate infection by bacteriophage Sf6. J Mol Biol 427:1964–1976.

Reyes-Robles T, Dillard RS, Cairns LS, Silva-Valenzuela CA, Housman M, Ali A, Wright ER, Camilli A. 2018. Vibrio cholerae Outer Membrane Vesicles Inhibit Bacteriophage Infection. J Bacteriol 200.

Steinbacher S, Baxa U, Miller S, Weintraub A, Seckler R, Huber R. 1996. Crystal structure of phage P22 tailspike protein complexed with Salmonella sp. O-antigen receptors. Proc Natl Acad Sci U S A 93:10584–10588.

Steinbacher S, Seckler R, Miller S, Steipe B, Huber R, Reinemer P. 1994. Crystal structure of P22 tailspike protein: interdigitated subunits in a thermostable trimer. Science 265:383–386.

Subramanian S, Parent KN, Doore SM. 2020. Ecology, Structure, and Evolution of Shigella Phages. Annu Rev Virol 7:121–141.

Sun Q, Knirel YA, Lan R, Wang J, Senchenkova SN, Shashkov AS, Wang Y, Wang Y, Luo X, Xu J. 2014. Dissemination and serotype modification potential of pSFxv_2, an O-antigen PEtN modification plasmid in Shigella flexneri. Glycobiology 24:305–313.

Tan D, Svenningsen SL, Middelboe M. 2015. Quorum Sensing Determines the Choice of Antiphage Defense Strategy in Vibrio anguillarum. mBio 6:e00627.

Teh MY, Furevi A, Widmalm G, Morona R. 2020. Influence of Shigella flexneri 2a O Antigen Acetylation on Its Bacteriophage Sf6 Receptor Activity and Bacterial Interaction with Human Cells. J Bacteriol 202.

Teh MY, Tran ENH, Morona R. 2022. Bacteriophage Sf6 host range mutant that infects Shigella flexneri serotype 2a2 strains. FEMS Microbiol Lett.

Tinney KR, Dover JA, Doore SM, Parent KN. 2022. Shigella viruses Sf22 and KRT47 require outer membrane protein C for infection. Biochim Biophys Acta Biomembr 1864:183920.

Trojet SN, Caumont-Sarcos A, Perrody E, Comeau AM, Krisch HM. 2011. The gp38 adhesins of the T4 superfamily: a complex modular determinant of the phage’s host specificity. Genome Biol Evol 3:674–686.

Tzipilevich E, Habusha M, Ben-Yehuda S. 2017. Acquisition of Phage Sensitivity by Bacteria through Exchange of Phage Receptors. Cell 168:186–199 e112.

van der Ley P, de Graaff P, Tommassen J. 1986. Shielding of Escherichia coli outer membrane proteins as receptors for bacteriophages and colicins by O-antigenic chains of lipopolysaccharide. J Bacteriol 168:449–451.

Wang C, Tu J, Liu J, Molineux IJ. 2019. Structural dynamics of bacteriophage P22 infection initiation revealed by cryo-electron tomography. Nat Microbiol 4:1049–1056.

West NP, Sansonetti P, Mounier J, Exley RM, Parsot C, Guadagnini S, Prevost MC, Prochnicka-Chalufour A, Delepierre M, Tanguy M, et al. 2005. Optimization of virulence functions through glucosylation of Shigella LPS. Science 307:1313–1317.

Zaidi MB, Estrada-Garcia T. 2014. Shigella: A Highly Virulent and Elusive Pathogen. Curr Trop Med Rep 1:81–87.

