## Supplementary Tables 1, 2, 3; Supplementary Figure 1 for "Host range expansion of *Shigella* phage Sf6 evolves through the dual roles of its tailspike"

Supplementary Table 1. List of Sf6 mutant isolates, earliest passage in which they arose, and total number of isolations

| Nucleotide change | Amino acid change | Earliest Isolation | Total Isolations |
|---|---|---|---|
| C14998G | A426G | 6 | 16 |
| C14998G/A15085T | A426G/N456I | 4 | 1 |
| C14998G/A15244C | A426G/N509T | 17 | 1 |
| C14998G/A15411G | A426G/T564R | 17 | 1 |
| A15048C | T443P | 6 | 1 |
| A15085T | N455I | 4 | 11 |
| A15085T/A15387T | N455I/N556Y | 4 | 1 |
| G15475A | G585D | 7 | 1 |
| T15486G | S589A | 11 | 1 |

Supplementary Table 2. Strains of bacteria used in this study

| | **Description** | **Reference** |
|---|---|---|
| **Bacteria** | | |
| PE577 | *S. flexneri* serotype Y (-:3,4) | (Morona, et al. 1994) |
| PE577/*ΔompA* | *ompA::kan*$^R$ | This study |
| PE577/*ΔompC* | *ompC::kan*$^R$ | This study |
| PE577/*ΔompAC* | *ΔompC/ompA::kan*$^R$ | This study |
| PE577/*ΔwaaL* | *waaL::kan*$^R$ | This study |
| CFS100 | *S. flexneri* serotype 2a2 (II:9,10) | (Marman, et al. 2014) |
| CFS100/*ΔompA* | *ompA::kan*$^R$ | This study |
| CFS100/*ΔompC* | *ompC::kan*$^R$ | This study |
| CFS100/*ΔompAC* | *ΔompC/ompA::kan*$^R$ | This study |
| CFS100/*ΔwaaL* | *waaL::kan*$^R$ | This study |
| CFS100/*ΔgtrII* | *gtrII::kan*$^R$ | This study |
| CFS100/*ΔgtrIIΔompA* | *ΔompA/gtrII::kan*$^R$ | This study |
| CFS100/*ΔgtrIIΔompC* | *ΔompC/gtrII::kan*$^R$ | This study |

Supplementary Table 3. Parameters of cryo-EM data collection and processing for Sf6 bound to CFS100 LPS

| | |
|---|---|
| Magnification | 53,000 |
| Voltage (keV) | 300 |
| Electron exposure (e–/Å2) | 33 |
| Defocus range (μm) | 0.8 – 3.0 |
| Pixel size (Å) | 0.816 |
| Symmetry imposed | C1 |
| Number of micrographs | 6241 |
| Final particle images (no.) | 18,964 |
| Map resolution (Å)<br>FSC threshold | 5.1<br>0.143 |

Supplementary Figure 1. Experimental evolution workflow

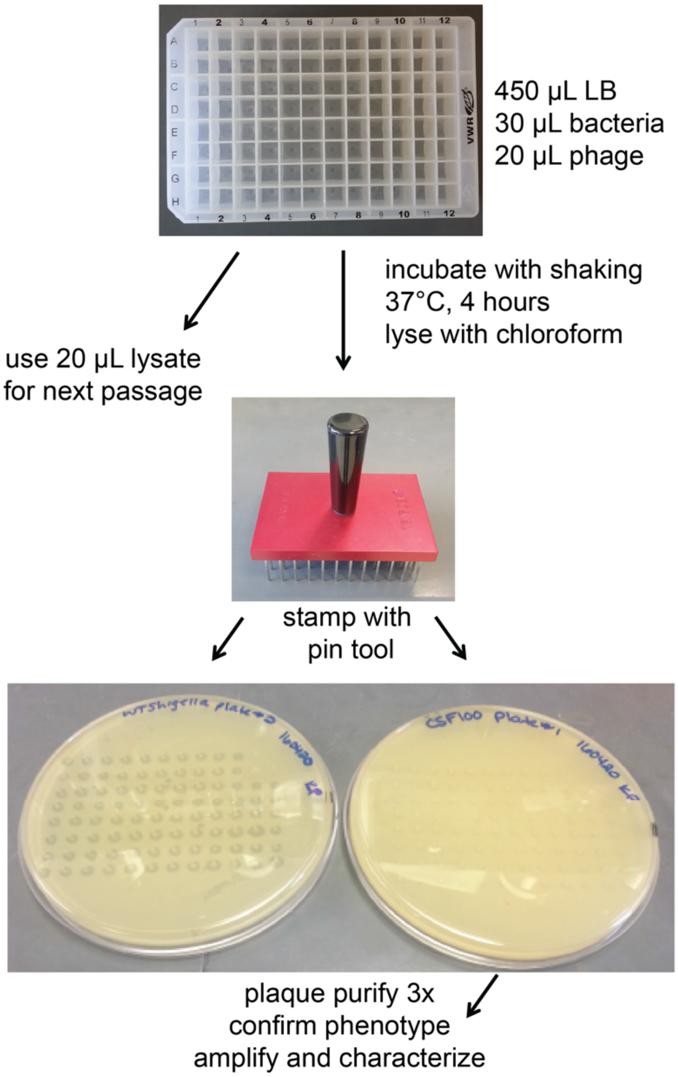
